# CASK hypomorph mice display cone photoreceptor dysfunction

**DOI:** 10.64898/2026.01.13.698659

**Authors:** Sheida Hashemi, Sara Y. Sabbagh, Khushi Talajia, James Fortenberry, Saba Tufail, Candice Turner, Konark Mukherjee

## Abstract

Variants in the X-linked gene *CASK* are associated with neurodevelopmental defects. Animal model studies have demonstrated that conditions such as cerebellar hypoplasia, microcephaly, and optic nerve hypoplasia (ONH) are related to loss-of-function (LOF) in the *CASK* gene. *CASK* variants are associated with multiple ocular conditions spanning both anterior and posterior segments of the eye including retinopathies. Both *Cask* heterozygous knockout (+/−) mice and *Cask* knock-in (KI) mice with reduced *Cask* expression have been shown to display ONH. *Cask* (+/−) mice displayed no defects in retinal structure or function. Here, we have systematically examined the *Cask* (KI) mice. Our results demonstrate that the anterior segment of the eye in *Cask* (KI) mice does not display any obvious phenotype. *Cask* (KI) mice however show a reduced visual acuity in optomotor response. The retina of *Cask* (KI) mice does not exhibit any major changes in their structure, vasculature, or gene expression pattern. We, however, uncovered a specific dysfunction of the cone receptor in *Cask* (KI) mice using electroretinogram (ERG). Mechanistically this dysfunction arises due to lowered levels of cone-specific opsin (opsin1mw) in *Cask* (KI) mice. To the best of our knowledge, this is the first description of retinal dysfunction in an animal model with *CASK* gene suppression. We infer that like ONH and cerebellar hypoplasia, retinopathy also may represent *CASK* LOF.

## Introduction

Mutations in the X-linked intellectual disability gene *CASK* (calcium/calmodulin-dependent serine protein kinase) have been associated with a condition known as MICPCH (microcephaly and pontocerebellar hypoplasia; OMIM) (1, 2). *CASK* is a multidomain protein composed, from the N- to the C-terminus, of a calcium/calmodulin-dependent kinase (CamK) domain, two LIN2/7 (L27) domains, a PSD95/Dlg1/ZO-1 (PDZ) domain, a Src-homology 3 (SH3) domain, and a guanylate kinase (GuK) domain (3). Numerous interactions of CASK have been described in the literature (4-8). The CaMK domain of CASK is an atypical kinase and may phosphorylate CASK-interacting synaptic proteins such as neurexin (9, 10). Although CASK is known to have a synaptic function, its wide tissue distribution suggests that it may have additional cellular or molecular functions that remain uninvestigated (3, 11).

Besides microcephaly and pontocerebellar hypoplasia, individuals with variants in the *CASK* gene may display global developmental delay, seizures, hypotonia, sensorineural deafness, and ocular abnormalities (12-14). Occasionally, defects in the heart and kidney have also been noted (15, 16). All *CASK*-linked manifestations exhibit variable penetrance and expressivity. Boys with missense mutations may exhibit intellectual disability, with or without nystagmus, in the absence of MICPCH (17). Seizures are present in approximately 40% of cases (18). Microcephaly and developmental delay may also manifest in the absence of pontocerebellar hypoplasia (19). These differences in manifestation are likely to depend on the nature of the mutation, the sex of the individual, as well as genetic background and other stochastic factors.

Many ocular phenotypes have been associated with *CASK* variants, including nystagmus, strabismus, megalocornea, stellate iris, glaucoma, cataract, persistent hyperplastic primary vitreous, retinopathies, optic nerve hypoplasia (ONH), and optic nerve atrophy (12, 13, 16, 17, 20). The role of CASK in eye development is therefore complex and multifaceted. How CASK influences overall eye development remains to be comprehensively investigated. *Cask* mutant mouse models recapitulate many of the known MICPCH phenotypes. *Cask* (+/-) mice display microcephaly, sporadic seizures, cerebellar hypoplasia, decreased body weight and scoliosis (21-23). *Cask* (+/-) mice were generated from a mouse line where the exon 1 of *Cask* gene is flanked by loxP sequences. In the 3’ adjacent intronic segment a neomycin resistance cassette was placed followed by a third loxP sequence. It was observed the floxed line which included the neomycin resistance cassette had a lower expression of CASK protein and is thus a hypomorph (24) (Figure 1A). Previous studies on animal models have demonstrated that *Cask* haploinsufficiency, as well as hypomorphism, may be associated with ONH. The overall structure and function of the retina however remain intact in the *Cask* (+/–) haploinsufficient mice (21, 25).

**Figure 1.**
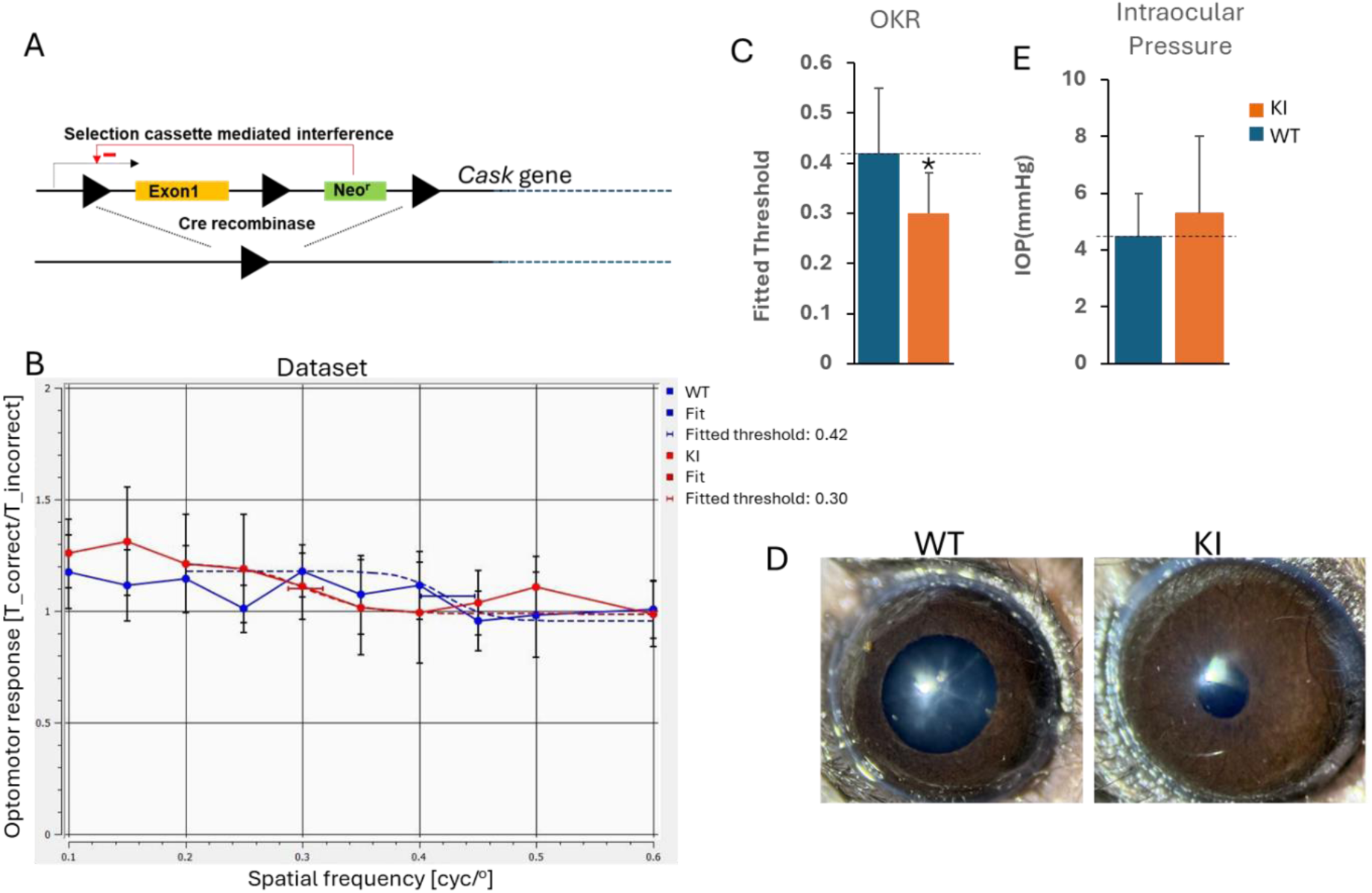
CASK KI mice have a visual deficit but a normally developed eye. A) Design of the CASK KI mouse line. B) OKR spatial frequency threshold assessment WT and CASK KI mice, The x-axis shows spatial frequency (cycles/degree), which represents the fineness of the visual gratings used in the OKR test. The y-axis displays the threshold OKR response, defined as the highest spatial frequency at which mice can reliably track the visual stimulus. WT mice demonstrate a higher mean OKR threshold (0.45 cycles/degree) compared to KI mice (0.3 cycles/degree; mean ± SEM; *p* < 0.05) C) Fitted threshold of indicated genotype plotted as mean±SEM, * indicates P<0.05, N=4. D) Representative slit lamp images from mice with indicated genotypes. E) Intraocular pressure plotted as mean±SEM from mice of indicated genotype. N=4.

Herein, we have investigated a *Cask* floxed knock-in (KI) mouse, which displays ONH (25). Overall, eye development in *Cask* KI mice appears unhampered. The structure and gene expression of the neuroretina, despite a reduced number of retinal ganglion cells (RGCs) (25), are not very different than those of the wild type mice. We observed, however, a specific reduction in the amplitude of the light-adapted a- and b-waves of the electroretinogram (ERG), indicating a selective cone dysfunction. We found that a reduced level of cone-specific opsin (opsin1mw) underlies this cone dysfunction. This is the first description of a retinopathy in an animal model of *CASK* mutation.

## Materials and Method

### Statement of ethics

All animal procedures were performed in accordance with the guidelines of the Institutional Animal Care and Use Committee (IACUC) of University of Alabama at Birmingham. Mice were maintained and genotyped as previously described (25). Animals used in the studies were between 12-14 months of both sexes. Animals were euthanized under photopic conditions.

### Optokinetic response (OKR)

Mice were briefly acclimated to the experimental arena before testing. Each animal underwent four trials to allow sufficient averaging of visual response curves. Platforms and contrast plates were selected according to coat color to optimize tracking. Optokinetic response (OKR) measurements were performed using the PhenoSys qOMR system (PhenoSys GmbH, Berlin, Germany). The arena comprised of four monitors arranged around a central platform, with infrared illumination adjusted for maximum contrast between the mouse and the background. Camera focus was preset, and images were captured at 30 frames per second (fps).

Visual stimuli consisted of sinusoidal gratings at varying spatial frequencies, with stimulus sequences randomized for each session to minimize order effects. Each trial presented a rotating pattern, with parameters specifying spatial frequency, contrast, speed, and duration. A 20-second gray interval was included between gratings to promote animal attention.

Real-time head-position tracking was achieved through automated image analysis, with experimenters able to adjust tracking parameters when required. All behavioral data including head-movement responses correlated with stimulus direction were recorded and exported for subsequent analysis. Behavioral responses were quantified as the ratio of correct to incorrect head movements relative to stimulus rotation. Data from multiple trials were plotted against spatial frequency, and sigmoid-curve fitting was applied to estimate acuity thresholds. Experimental sessions were scheduled in alignment with animal circadian rhythms and lasted no more than 30 minutes per animal.

### Intraocular Pressure Measurement and Slit-lamp Examination

Mice were anesthetized via intraperitoneal injection of ketamine and xylazine (50 mg/kg and 5 mg/kg, respectively) and placed on a heated platform to preserve body temperature. Intraocular pressure (IOP) was measured using the iCare IC100, a handheld rebound tonometer. Although the device is designed for human use, the1.8 mm disposable probe is sufficiently small to allow accurate IOP measurements in mice. While under anesthesia, slit lamp biomicroscopy was performed using the Keeler PSL Classic Portable Slit Lamp. High-resolution images of the cornea and iris were captured using an iPhone 16 Pro Max mounted to the slit lamp ocular via the Keeler Eye Mobile Smartphone Adapter.

### Electroretinogram

Mice underwent 2 hours of dark adaptation before being anesthetized with a ketamine–xylazine mixture (50 mg/kg and 5 mg/kg, respectively). Corneal anesthesia and pupil dilation were performed prior to recording. Electroretinogram (ERG) signals were captured using a contact lens electrode coupled with 2.5% methylcellulose, while a gold-wire reference electrode was positioned on the contralateral (unstimulated) eye. ERGs were recorded using a custom system that delivered light stimuli (505nm) via fiber optics from either a 100-W halogen lamp or a high-intensity LED source. Dark-adapted responses were elicited by brief flashes spanning intensities from 0.622 to 6.316 log photons/μm². After a 3-minute exposure to rod-saturating light for light adaptation, responses to flashes at 6.955 log photons/μm² were recorded. The amplitude of the a-wave was measured from the baseline to its trough, while the b-wave amplitude was calculated from the a-wave trough to the b-wave peak. Implicit times were determined using stimuli at the highest intensities presented.

### Optical Coherence Tomography (OCT)

Mice were anesthetized via intraperitoneal injection of ketamine and xylazine (50 mg/kg and 5 mg/kg, respectively) and maintained on a heated platform to preserve body temperature. Pupils were dilated using phenylephrine (10%; Lifestar Pharma LLC, C4X001) and tropicamide (0.1%; Somerset Therapeutic LLC, A230527) eye drops administered at least 15 minutes before imaging. Corneal hydration was maintained with ophthalmic gel throughout the procedure.

Retinal imaging was performed using a Bioptigen Model 840 Envisu Class-R high-resolution spectral-domain optical coherence tomography (SD-OCT) system (Bioptigen/Leica, Inc., Durham, NC, USA) equipped with a mouse-specific contact lens. The system included a base unit, an animal imaging mount with multi-axis translational stages, and a probe aligned to the eye using a built-in bite bar and nose band to stabilize head position.

Following anesthesia and positioning, the eye was aligned centrally under the OCT probe using micromanipulators to ensure precise focus on the optic nerve head and surrounding retina. Multiple scan types were acquired, including radial, rectangular, and annular scans, each defined by parameters such as diameter, number of A-scans per B-scan, and total volume frames.

OCT images and volumetric data were collected and stored using Bioptigen proprietary software. Each scan session was saved with unique identifiers corresponding to the animal and eye examined. Retinal layer thickness measurements focused on layers containing retinal cells, such as the retinal nerve fiber layer (RNFL) and ganglion cell layer (GCL). Automated segmentation software provided initial layer delineations, and measurements were refined using customized image-processing macros when necessary. Multiple measurement points were analyzed per scan to improve accuracy.

After imaging, mice received additional ophthalmic gel to prevent corneal desiccation and were monitored until full recovery from anesthesia on a warmed surface.

### RNA Sequencing and Data Analysis

RNA was extracted from whole dissected retina following the manufacturer’s instruction using the Aurum total RNA fatty and fibrous tissue kit^TM^ (Biorad). RNA quality and integrity were assessed using an Agilent Bioanalyzer. Sequencing libraries were prepared using the NEB Ultra II Directional RNA Library Prep Kit and sequenced on an Illumina NovaSeq 6000 platform. Raw reads were quality-checked and trimmed using *Trim Galore* along with *Cutadapt*. Clean reads were aligned to the *Mus musculus* reference genome (GRCm39/mm39) using *STAR* aligner. Gene-level counts were generated with *HTSEQ*, and differential expression analysis was performed using *DESeq2*.

### Immunohistochemistry

Mouse whole retinas were fixed overnight in 4% paraformaldehyde (PFA) at 4°C, followed by three washes in phosphate-buffered saline (PBS) for 10 minutes each, and then cryoprotected overnight at 4°C in 30% sucrose. Samples were permeabilized and blocked overnight at 4°C in blocking buffer (10% horse serum (Lonza, Cat# 14-403F) prepared in PBS with 0.1% Triton X-100 (PBST)).

Primary antibodies, cone arrestin (Sigma-Aldrich, Cat# AB15282), GNAT2, BSA-free (Novus Biologicals, Cat# NBP3-13378), and Opsin 1 (medium-wave) (Novus Biologicals, Cat# NB110-74730) were each diluted 1:200 in blocking buffer and incubated overnight at 4°C. After three washes in PBST for 30 minutes each, retinas were incubated in the dark with Alexa Fluor 633 goat anti-rabbit IgG secondary antibody (Thermo Fisher Scientific, if applicable) diluted 1:500 in blocking buffer for 3 hours at room temperature. Final washes were performed before cryosectioning and mounting the retinas on glass slides with antifade mounting medium for confocal imaging. Imaging was done using Keyence digital microscope and Zeiss LSM 710 inverted confocal microscope.

### Fluorescein angiography

Fundoscopy in mice was performed using the Phoenix MICRON® IV retinal microscope, enabling high-resolution imaging of the fundus, optic disc, and retinal vasculature. Mice were anesthetized by intraperitoneal injection of ketamine (50 mg/kg) and xylazine (5 mg/kg), and their pupils were dilated with 1% tropicamide and 2.5% phenylephrine. To preserve corneal moisture and ensure optical clarity, a sterile ophthalmic lubricant was applied and, when necessary, a miniature contact lens was gently placed on each eye. Mice received 100 µl of 1% fluorescein (Fluorescite^®^) intraperitoneally. The animals were maintained on a thermostatically controlled heating pad throughout the procedure to stabilize body temperature. Retinal images were acquired by methodically adjusting the microscope’s optical alignment until the central retina and peripheral vessels were in sharp focus, and Phoenix Discover software was used to capture and process images in real time. Lubricant was reapplied as needed to maintain corneal transparency, and light exposure was minimized to prevent retinal phototoxicity. Once imaging was complete, mice were placed in a recovery chamber and monitored until full locomotor activity was restored. This protocol incorporates standard fundus imaging steps and integrates Phoenix MICRON IV’s system-specific features for optimal image resolution and animal safety.

## Results

### Normal eye development and reduced visual acuity in CASK hypomorph mice

Previously, it has been reported that *Cask* floxed knock-in mice have reduced CASK expression in their nervous system (24). These mice display optic nerve hypoplasia (ONH) with a reduction in the number of retinal ganglion cells (RGCs) and thinning of the optic nerve, as well as of individual RGC axons that course through the optic nerve (25). To date, no visual functional study has been carried out on these animals. Here, we performed an optokinetic response (OKR) test. The OKR is an evolutionarily conserved visual response and provides objectivity in measurement due to its reflexive nature (26). The OKR signals are carried by axons of specialized RGCs, called ON direction-selective RGCs, to the midbrain. It was therefore expected that the OKR would be altered in the *Cask* knock-in (KI) mice due to ONH and loss of RGC axons to the midbrain. Our OKR result show that both groups KI (red) and WT (blue) display a ratio of ∼1–1.3, and their curves overlap with no obvious separation of the groups at specific spatial frequencies. This suggests broadly similar optomotor performance across the tested spatial frequency range for both WT and KI mice. The dashed lines are ‘fitted psychometric/threshold functions’ for WT and KI mice, the fitted thresholds for the WT mice are 0.42 cyc/° whereas for the KI mice it is 0.30 cyc/°. Our experiment thus indicates that the KI mice lose reliable tracking at a coarser (lower-frequency) pattern than WT, implying reduced spatial resolution/acuity in the optomotor task (Figure 1B,C). Although the reduced acuity in these mice could stem from ONH that has been already described, CASK variants are associated with other ocular conditions that may affect vision. We therefore systematically examined the eyes of the *Cask* KI mice.

Slit-lamp examination of the *Cask* KI mice revealed that, overall, the corneas and irises of the *Cask* KI mice did not display any developmental deviations from those of the wild-type controls (Figure 1D). In some cases, we observed lenticular clouding, which resulted from ketamine anesthesia. Since subjects with CASK variants may also develop juvenile glaucoma, we examined the intraocular pressure (IOP) in the *Cask* KI mice. Similar to the iris and cornea, the IOP of the KI mice was indistinguishable from that of the wild-type controls (Figure 1E). Overall, our data indicates that anterior segment of the eye development is normal in the *Cask* KI mice.

### No structural or gene expression changes in the retina of CASK hypomorph mice

*Cask* (+/–) mice display ONH and a reduction in the number of retinal ganglion cells (RGCs); however, the retinae of *Cask* (+/–) mice have normal development and anatomy (21, 25). The *Cask* KI mice also displayed ONH and a reduction in the number of RGCs (25), but the dissected retinae of the *Cask* KI mice appeared identical in size and shape to those of the wild-type mice, indicating that, similar to the *Cask* (+/–) mice, the pathology in the *Cask* KI mice may be limited to the ganglion cell layer (GCL) (Figure 2A). To further examine retinal structure, we performed optical coherence tomography (OCT). Our data indicates that the retinal anatomy of the *Cask* KI mice is within the normal range and that the thickness of individual retinal layers remains unchanged (Figure 2B,C). Strikingly, although *Cask* KI mice have ONH, we did not observe a change in retinal nerve fiber layer (RNFL) thickness. One possible explanation is that the RNFL in mice is very thin and near the resolution limits of OCT (27). Overall, our data indicates that the reduction of CASK has no observable effect on retinal formation. ONH may be associated with vascular tortuosity (28). Furthermore, CASK has been postulated to have a role in the endothelium (29), and CASK variants have been associated with cardiovascular defects (30). We therefore performed fluorescein angiography in the *Cask* KI mice and wild-type controls. In both cases, we observed clear lenses, indicating that, unlike some reported clinical cases involving CASK variants, the *Cask* KI mice did not display cataracts. Overall, the vasculature in the *Cask* KI mice was indistinguishable from that of the wild-type controls. We quantified the diameter of the major vessels, their branching as well as total number of pixels included in retinal vascularization. All values were identical between the wildtype controls and the *Cask* KI mice (Figure 2 D, E, F and G).

**Figure 2.**
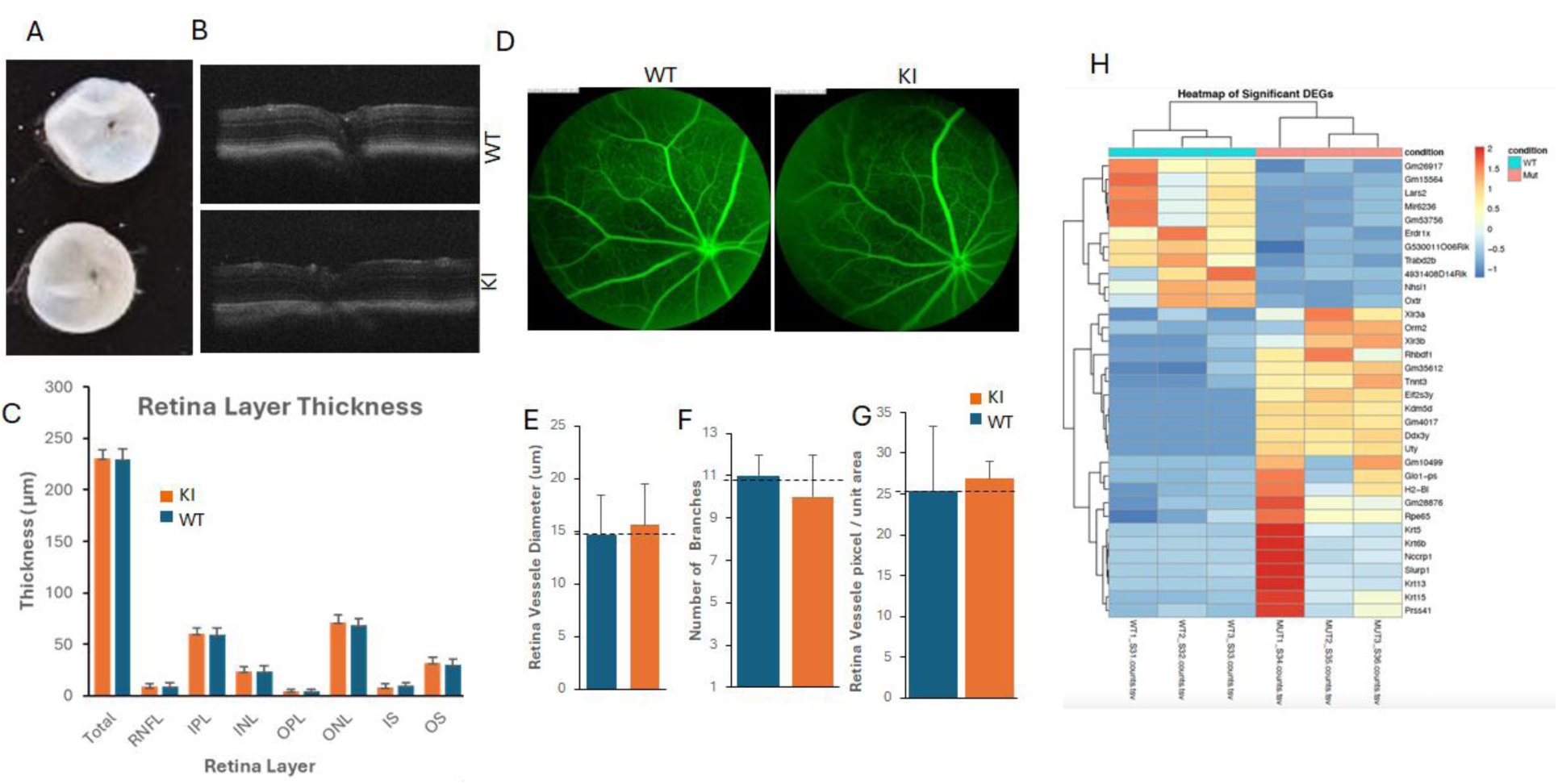
Retina of CASK KI has a normal retinal structure and gene expression. A) Representative images of dissected whole retina from mice of indicated genotype. B) Representative images of OCT from indicate mice. C) Quantitation of OCT, results plotted as mean and SEM. RNFL= retinal nerve fiber layer, IPL= inner plexiform layer, INL= inner nuclear layer, OPL= outer plexiform layer, ONL= outer nuclear layer, IS= inner segment, OS= outer segment, RPE= retinal pigment epithelium. D) Representative images from fluorescein angiography of retina from mouse of indicated genotype. E, F, G) Quantitation of indicated parameters from the fluorescein angiographs. Results are plotted as mean and SEM, n=3 mice. H) Heatmap obtained from bulk RNA-seq from retina of WT and CASK KI mice n=3 mice in each group.

We therefore next examined potential changes in gene expression in the *Cask* KI mice using RNA sequencing (RNA-seq). Although CASK has a purported role in gene transcription (7), RNA sequencing of the brains of *Cask* (+/–) mice has previously failed to reveal any major changes, except those related to extracellular matrix genes, which are expected in degenerative conditions. Most of the molecular alterations in *Cask* (+/–) mice are post-translational in nature (22). In our current study, we examined whether the retinae of the *Cask* KI mice display any gene expression changes indicative of aberrant retinal biochemistry. Similar to the *Cask* (+/–) mice, we did not detect major changes in transcript levels in the retinae of the *Cask* KI mice compared to wild-type controls. We analyzed the results as described in the *Methods* section. More than 26,000 RNAs were sequenced per sample. Using log₂ fold change analysis, we identified only 64 differentially expressed genes (DEGs): 25 were downregulated, and 39 were upregulated (Figure 2H). Due to the small number of DEGs, further enrichment analysis was not feasible. Gene ontology classification indicated that many of the genes with altered expression are localized to the plasma membrane and intermediate filaments and may be involved in signaling and cytoskeletal organization (not shown). Overall, our data indicates that a decrease in CASK levels does not alter the developmental trajectory of the retina and has only a limited effect on gene expression.

### CASK hypomorph mice display a specific reduction in light-adapted ERG response due to lowered levels of cone opsin

We next examined the retinal function of the *Cask* KI mice using ERG, as described in the *Methods* section. Previous studies have failed to detect any deficits in the ERG of *Cask* (+/–) mice (25), which exhibit mosaic expression of CASK due to its X-linkage. ERG waveforms are well-established indicators of outer retinal neural activity: the a-wave represents photoreceptor responses, whereas the b-wave corresponds to the activity of depolarizing bipolar cells. We performed ERG recordings under both dark-adapted (scotopic) and light-adapted (photopic) conditions.

Because mice are nocturnal, we first examined the scotopic ERG responses. Under scotopic conditions, the a- and b-wave responses appeared similar between *Cask* KI and wild-type mice at all retinal illuminances (Figure 3A, B and C). Strikingly, under photopic conditions, both the a-and b-waves were markedly diminished in the *Cask* KI mice across all light intensities (Figure 3D, E and F). Taken together, these findings indicate that cone-driven responses at 505 nm are specifically reduced in *Cask* KI mice. We therefore next examined molecules involved in cone phototransduction.

**Figure 3.**
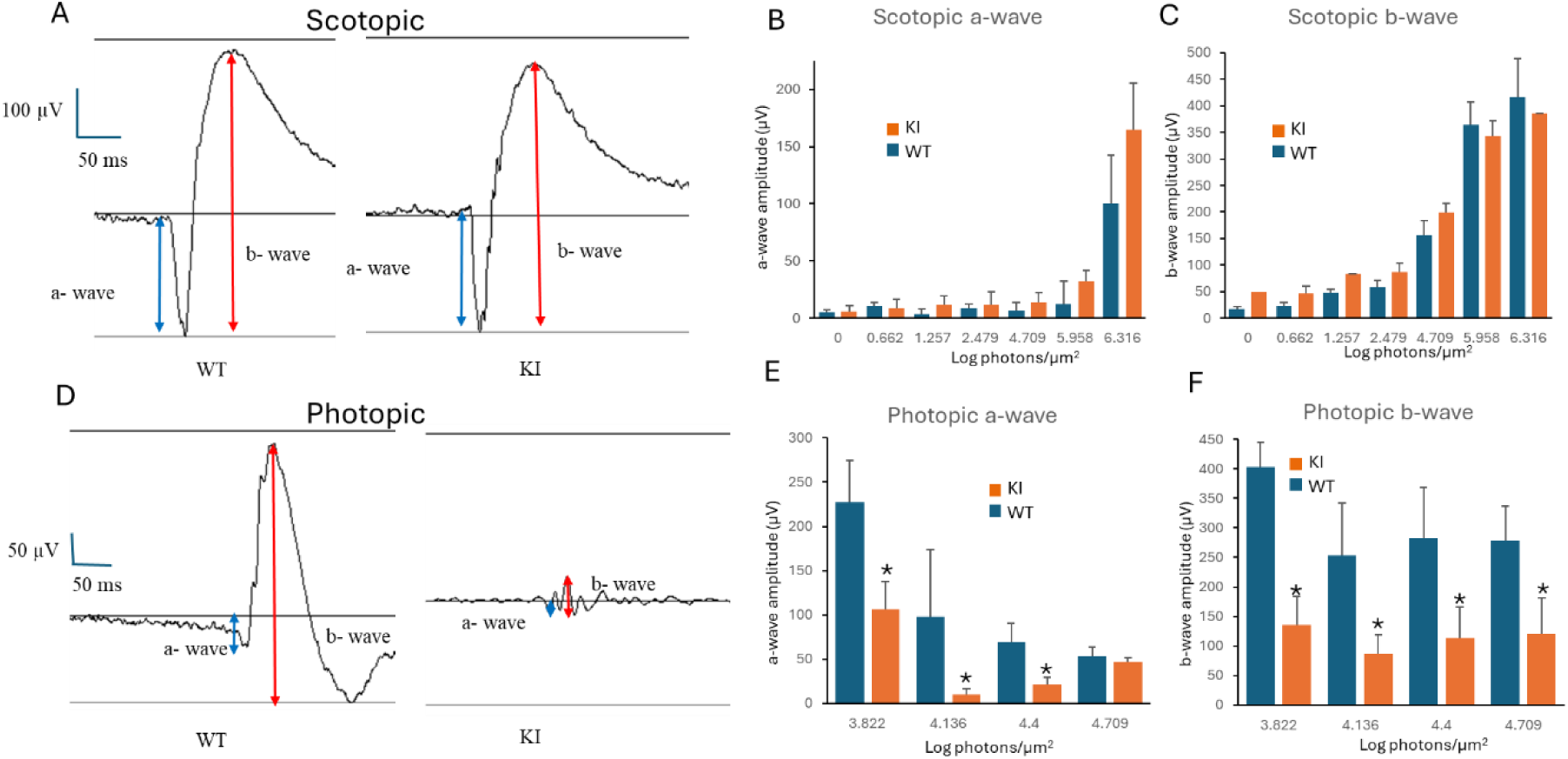
Retina of C*ask* KI has a reduced cone-driven response. Electroretinogram (ERG) responses in wildtype controls and CASK KI mice. (A, D) Representative ERG waveforms showing a- and b-wave responses under scotopic and photopic conditions in WT and KI mice. (B, C) Quantification of scotopic a-wave and b-wave amplitudes across increasing light intensities shows no significant difference between groups. (E, F) Photopic a-wave and b-wave amplitude comparisons reveal significantly reduced responses in KI mice relative to WT at multiple light intensities (mean ± SEM; * indicates P<0.05); N=4.

Cones represent a minor photoreceptor subtype in mammals; in mice, they constitute only ∼3% of the total number of photoreceptors (31). Phototransduction involves light-mediated isomerization of photopsins, which triggers signaling through the G-protein transducin, leading to membrane hyperpolarization due to ion channel closure (32). This process is terminated by specific β-arrestins. We therefore quantified cone receptor–specific levels of β-arrestin, transducin, and opsin using immunolabeling experiments. Figure 5 depicts these labeling results; we did not observe any significant differences in the labeling pattern or intensity of either β-arrestin-C or transducin (GNAT2) between *Cask* KI and wild-type mice (Figure 4A,B, D and E). These results suggest that there is no overall decrease in the number of cone photoreceptors in *Cask* KI mice and that these cells express β-arrestin and transducin at normal levels.

**Figure 4.**
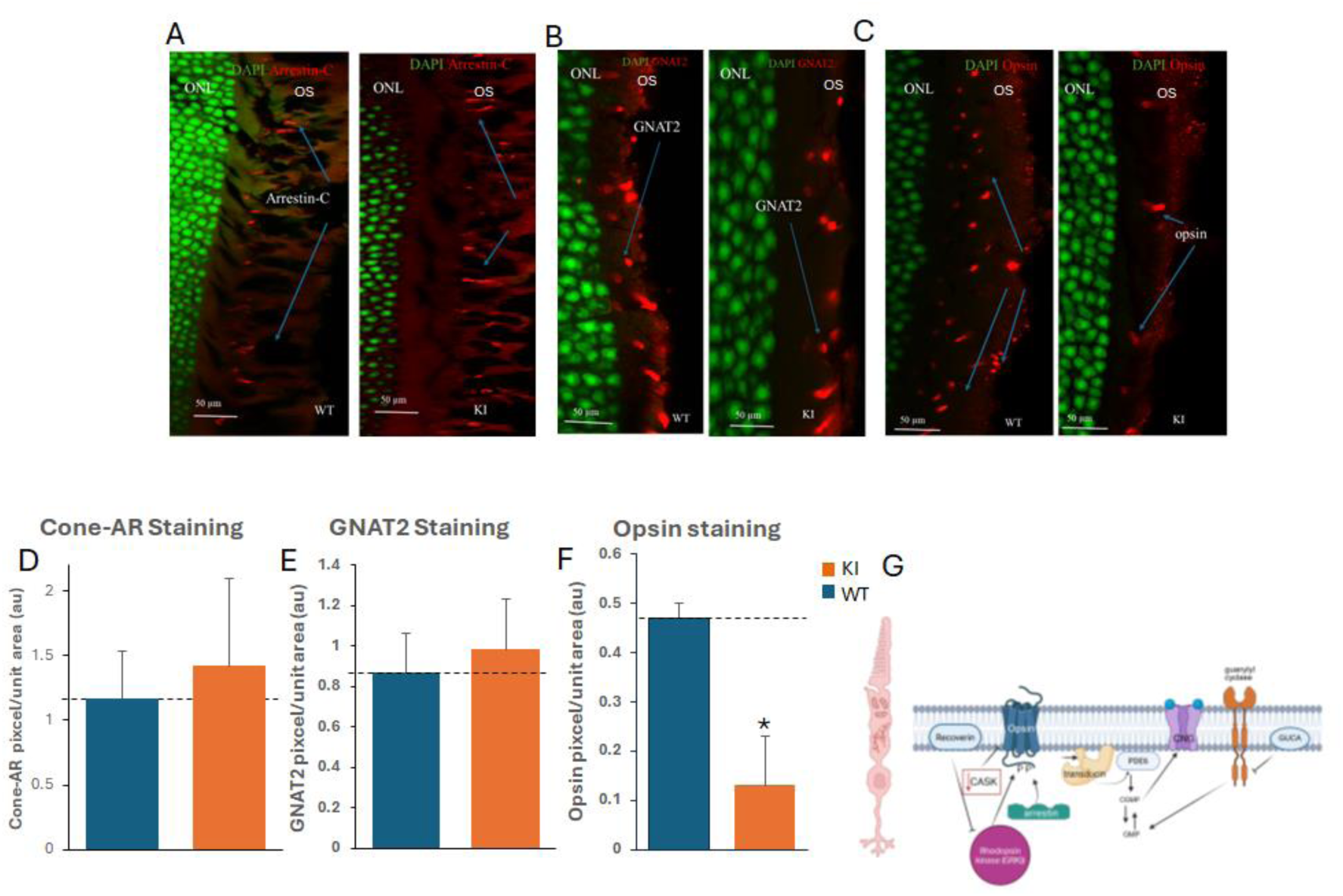
Reduction of opsin 1mw levels in CASK KI mice. A, B and C) Representative images of outer retina from mice of indicated genotype immunolabeled for arrestin-C, GNAT2 and opsin1mw. The nucleus in ONL (outer nuclear layer) is labeled with DAPI (green), OS is outer segment. Each of the mentioned antigens is labeled with red color. Scale bars are provided in each image. D, E and F) Quantitation of immunolabeling experiments plotted as mean and SEM. * indicates P<0.05, N=3 mice per group. G) A model of phototransduction in cone cells. Decrease in CASK is associated with a decrease in opsin amounts. GUCA is guanylate cyclase activator, PDE6 is phosphodiesterase 6 and CNG is cyclic nucleotide-gated channels.

Mice are dichromatic and possess three types of cones based on the opsin expressed: true S-cones (UV-sensitive), true M-cones (green-sensitive), and dual cones. Cone distribution is nonuniform across the mouse retina. In pigmented strains such as C57BL/6, the ventral retina is rich in true S-cones (a minor cone subpopulation) and dual cones, whereas the dorsal retina is sparse in S-cones and predominantly contains M-cones (33). Therefore, to quantify cone receptor–specific opsin, we focused on opsin1 (M-type). This decision was also because the wavelength of the light source used in our ERG had wavelength very close to opsin1 mw sensitivity optima (∼508nm). Our results indicate that opsin1 mw levels were reduced approximately fivefold in the *Cask* KI retina compared to wild-type controls (Figure 4C and F), accounting for the reduction in the photopic ERG response. Overall, our data indicates that *Cask* KI mice exhibit specific cone receptor dysfunction caused by decreased opsin levels, without changes in the absolute number of cone photoreceptors (Figure 4G).

## Discussion

Incomplete penetrance and variable expressivity are common features of many monogenic rare disorders (34). These non-Mendelian phenomena are not well understood; however, several factors, such as environmental influences, genetic background, and epigenetic regulation may affect phenotypic expression (35). Furthermore, the development of complex systems, such as embryogenesis, is likely to involve multiple stochastic processes. Variants in the X-linked gene *CASK* may present with manifestations ranging from asymptomatic cases to profound encephalopathy resulting in lethality. Many of these phenotypic differences can be explained by the nature of the mutation and the sex of the individual. However, phenotypic penetrance, even with the same mutation in the same sex, is often incomplete, complicating genotype–phenotype correlations in *CASK*-related disorders. It is possible to find the same mutation that may or may not be associated with manifestations such as cerebellar hypoplasia (36, 37), optic nerve hypoplasia (19), optic nerve atrophy, or nystagmus (36, 37). In particular, individuals with *CASK* variants can present with a variety of ocular manifestations involving both the anterior and posterior segments. In one series, 5 of 13 subjects displayed defects on electroretinography (ERG) (12). Retinopathy has been reported in individuals with *CASK* variants, often accompanied by optic nerve pathology. Typically, such retinopathy may affect the rod responses (37).

Our investigation of *Cask* (+/−) mice indicated that optic nerve hypoplasia (ONH) may result directly from *Cask* deficiency in C57BL/6 mice. These mice displayed a thin optic nerve and a postnatal reduction in retinal ganglion cell (RGC) numbers with complete penetrance. Yet, the mechanism underlying RGC reduction is not straightforward. In fact, complete deletion of *Cask* in approximately 90% of RGCs did not prevent their development or affect survival. Strikingly, apart from the reduction in RGC number, no perturbation was noted in the structure or function of the retina in *Cask* (+/−) mice. One possible explanation is that, due to X-linkage, *Cask* (+/−) mice are mosaic rather than hypomorphic. It is possible that a normal level of CASK in 50% of retinal cells is sufficient for physiological responses in mice ERG. The *Cask* KI mice have been reported to be hypomorphic and to display ONH and a reduction in RGC numbers. We therefore systematically phenotyped *Cask* KI mice ophthalmologically. In addition to a known reduction in RGC number, and a diminished visual acuity in optokinetic responses (OKR), the only defect we observed was a reduction in cone-driven responses due to decreased opsin levels. The reduction in the visual acuity may relate either to the ONH or cone dysfunction.

To the best of our knowledge, this is the first demonstration of retinopathy in a *CASK*-linked animal model. Yet, the result was an unexpected one, as individuals with *CASK* variants have not been reported to display cone-specific defects or color blindness. The variations in retinal phenotype associated with *CASK* deficiency in humans may arise from differences in genetic background, whereas the cross-species differences observed here may reflect underlying biological divergence as observed in multiple studies of CASK orthologs (24, 38, 39). Unlike mice, humans are diurnal and exhibit higher visual function during the daytime (photopic). In humans, cones are densely packed in the fovea, where their concentration is much higher than that of rods, and they are crucial for high-acuity vision (40). Cone receptor dysfunction can give rise to color blindness in humans (achromatopsia). Besides the loss of color vision, cone dysfunction also leads to photophobia, poor visual acuity, and nystagmus. Achromatopsia is typically monogenic and results from mutations in subunits of the cyclic nucleotide-gated channel and other molecules involved in phototransduction in cone cells (41).

Evidence indicates that mice, although dichromatic, can distinguish between colors (42). Here, we investigated the molecular basis of decreased cone activity in *Cask* KI mice. We examined three molecules: cone β-arrestin, which inactivates opsin; opsin1mw, due to its wider distribution in the retina; and transducin. Our data suggests that a specific decrease in cones expressing opsin1mw may account for the reduced cone-driven responses. Unfortunately, our bulk RNA-seq data failed to reveal any specific transcriptional signature linking retinal signaling to reduced CASK expression that could explain the lower amounts of opsin1mw.

Multiple observations to date have identified *CASK*-deficiency phenotypes that specifically affect certain cell types. For example, in the cerebellum, loss of CASK primarily leads to the death of granule cells but not Purkinje cells (19, 43). In the retina, we observe distinct effects of reduced CASK in different cell types: while there is a slight decrease in the number of RGCs, no apparent changes occur in bipolar cells, amacrine cells, or rod receptors. Cone receptors, however, show a specific loss of opsin1mw expression. The mechanism underlying this phenomenon remains unclear. We considered the possibility that the neomycin resistance cassette in the *Cask* gene might suppress opsin1mw gene expression, as both genes are located on the X chromosome; however, the genomic distance between them makes this explanation unlikely.

CASK protein interactions may be cell-type specific and modulated by alternative splicing (19, 44). Therefore, CASK likely performs multiple, distinct functions across cell types. It is plausible that CASK has a specific, yet unidentified, role in mouse cone receptors. Future studies are required to elucidate the role of CASK in cone receptor biology. To the best of our knowledge, our study is the first to report CASK-related retinopathy in an animal model, and the *Cask* KI mouse may serve as a valuable model for studying cone receptor dysfunction. Our data suggest that the retinopathies observed with CASK variants may be a direct result of *CASK* loss-of-function.

## Acknowledgment, authorship and funding.

SH, KT, SYS, CT, JF performed experiments, SH, ST, and KM analyzed and interpreted the data, KM conceptualized the project, SH and KM wrote the manuscript.

We thank Michael A Fox (UMASS Amherst) and Alecia Cross (UAB) for critically reading the manuscript. We thank VSRC core grant NIH P30 EY003039 at UAB. The study was done with support of R01EY033391. KM is supported by R01EY033391 and R01EY033141 from the National Eye Institute and Angelina CASK Neurological Research Foundation.

## Notes

### Competing Interest Statement

The authors have declared no competing interest.

